# GlyLES: Grammar-based Parsing of Glycans from IUPAC-condensed to SMILES

**DOI:** 10.1101/2022.11.10.515921

**Authors:** Roman Joeres, Daniel Bojar, Olga V. Kalinina

## Abstract

Glycans are important polysaccharides on cellular surfaces that are bound to glycoproteins and glycolipids. These are one of the most common post-translational modifications of proteins in eukaryotic cells. They play important roles in protein folding, cell-cell interactions, and other extracellular processes. Changes in glycan structures may influence the course of different diseases, such as infections or cancer.

Glycans are commonly represented using the IUPAC-condensed notation. IUPAC-condensed is a textual representation of the Symbol Nomenclature for Glycans (SNFG) that assigns a colored, geometrical shape to the main monosaccharides. These symbols are then connected in tree-like structures, visualizing the glycan structure on a polymeric level. Yet for a representation on the atomic level, notations such as SMILES should be used. To our knowledge, there is no easy-to-use, general, open-source, and offline tool to convert the IUPAC-condensed notation to SMILES. Here, we present the open-access Python package GlyLES for the generalizable generation of SMILES representations out of IUPAC-condensed representations. GlyLES uses a grammar to read in the monomer tree from the IUPAC-condensed notation. From this tree, the tool can compute atomic structures of each monomer based on their IUPAC-condensed descriptions. In the last step, it merges all monomers into the atomic structure of a glycan in the SMILES notation.

GlyLES is the first package that allows conversion between IUPAC-condensed notations of glycans and SMILES strings. This may have multiple applications, including straightforward visualization, substructure search, molecular modelling and docking, and a new featurization strategy for machine-learning algorithms. GlyLES is available at https://github.com/kalininalab/GlyLES.

## Introduction

### Glycans

In all kingdoms of life, cells, proteins, lipids, and other biological entities are often wrapped in chains of complex carbohydrates, known as glycans. Assembled by dedicated enzymes, glycosyltransferases, glycans form branched sequences from a pool of monosaccharides that is to some degree specific to each species. The specific glycan sequence that is attached to a macromolecule modulates its properties, such as structure, stability, or function. This fact, together with glycan’s diversity, makes glycans key in modulating functionality in most fundamental physiological processes [1]. For example, the lack of core fucosylation of the glycan attached to the antibody leads to increased antibody potency [2], or upregulation of sialyl-Lewis X structures facilitating the interaction with selectin proteins in cancer metastasis [3].

In fact, the modulation of protein binding is in some ways most sensitive to subtle changes in glycan chemistry. Glycan-binding proteins, or lectins, have evolved to recognize the exact shape and chemistry of particular glycan substructures or motifs. Witnessed naturally in viral infection, for example in the form of interaction of viral proteins with host glycan receptors [4], or the innate immune system [5], this property is also widely used in the laboratory and clinic, for enriching, visualizing, and targeting cells or proteins with characteristic glycan structures [6, 7]. Changes in glycan chemistry have also been observed in historic evolution (e.g., loss of functional groups such as the N-glycolyl group in humans [8]), as well as in contemporaneous evolution, in the form of the arms race in glycan-mediated host-microbe interactions [9, 10].

The structure of a glycan can be very complex due to many monosaccharides (~ 60), extraordinarily many branching structures, including different linkages, and the countless number of functional groups that can be attached to any monomer [11], particularly in bacteria.

Glycans are traditionally represented using so-called *IUPAC-condensed notation* which is a textual representation of the SNFG images [12]. An example of an SNFG visualization is shown in Figure 2a. It differs from the standard IUPAC, as illustrated by the example of mannose. In IUPAC, mannose has the description

~~~
(3S,4S,5S,6R)-6-(hydroxymethyl)oxane-2,3,4,5-tetrol,
~~~

whereas the IUPAC-condensed name is `Man`.

While this type of token-based notation has numerous advantages, including human-readability and compactness, chemical similarities between monosaccharides and substructures may occasionally be obscured. However, such similarities can substantially influence biochemical properties of the resulting glycan. Examples include the binding specificity of the glycan-binding protein WGA [13]. In IUPAC-condensed, its binding specificity (GlcNAc, GalNAc, Neu5Ac, MurNAc) seems overly broad. Yet, on a chemical level, WGA specifically binds to N-acetyl moieties shared by these monosaccharides. Other examples include lectins, such as MAL-I binding to negatively charged glycans [13], yielded by both sulfate groups or sialic acids. Thus, an increased resolution in glycan notation can provide more general biological insights into biological processes.

### SMILES

SMILES (Simplified Molecular-Input Line-Entry System) is a widely accepted standard for representing any chemical molecule as an ASCII string, in which non-hydrogen (heavy) atoms are encoded with their chemical symbols and a grammar is introduced for representing covalent bonds of different nature [14]. Whereas there are tools for inter-conversion between the standard IUPAC and SMILES, there are no such tools for the IUPAC-condensed notation of glycans. In this work, we aim to bridge this gap by introducing GlyLES, a Python-based open-source package, that given an IUPAC-condensed string for a glycan outputs the corresponding SMILES string.

### IUPAC-condensed notation of glycans

The IUPAC-condensed notation of glycans is the most commonly used notation in publications, databases, and other sources. Its wide usage is because of the compact descriptions of glycans and the human-readability of the notation [15].

This notation is a specialized form of the general IUPAC notation that is used to describe other organic molecules in a standardized way [16]. It is specifically suited for glycans and their variability in structure and attached functional groups in a compact form.

For simplicity, from now on we will refer to the IUPAC-condensed notation using the term “IUPAC” as the original IUPAC notation does not play a role in the presented work.

Glycans have a complex, yet regular, structure, especially when described using the IUPAC notation. For example,

~~~
Man(a1-4)Man(a1-4)Man(a1-4)Man
~~~

is a simple chain of four mannopyranose monosaccharides connected with 1-4 alpha-O-glycosidic bonds. Important to note is that the root of the glycan tree is always the last monosaccharide in an IUPAC formula. The chain grows from the end by prepending elements like `Man(a1-4)`, which corresponds to adding a new mannose to a leaf-monomer of the glycan. The branching of the trees is described by introducing the new branch into square brackets. For example,

~~~
Man(a1-4)Man(a1-3)[Man(a1-4)Man(a1-4)]Man(a1-4)Man
~~~

is a tree of mannopyranoses that has two monosaccharides before splitting into two chains with again two mannopyranoses each. The branching is put right in front of the monomer, where the root of the side branch is bound to the main strand. This can be extended to three chains of two monosaccharides, each bound to a single mannopyranose:

~~~
Man(a1-4)Man(a1-2)[Man(a1-4)Man(a1-3)][Man(a1-4)Man(a1-4)]Man.
~~~

Modifications in IUPAC are directly annotated at the changed monosaccharide. The annotation is often done by first naming the position of the carbon atom where a functional group is attached. Then, an abbreviation for this group is provided. So, for example, `Man3S` describes a mannose with a sulfur group attached to the third carbon atom. The number can also be dropped in case of a functional group attached to a standard position, as in `GalNAc` where an acetamide group is attached to the second carbon atom of galactopyranose. There are other possibilities to denote a functional group, but they are not addressed in this paper.

Modifications of glycans considered here include added sulfur groups (`Man3S`), added phosphate groups (`Man3P`), added amine groups (`ManN`). In total, GlyLES can process 59 monosaccharides (Supplementary Table S1) and 126 functional groups (Supplementary Table S2). Functional groups can be attached to any oxygen, nitrogen, or a carbon atom in the monosaccharide (e.g., `Man4S, Man4P`, or `ManNS`), and they can be combined in any way (e.g., `Man3S4P`). The only restriction is that one cannot attach two modifications to the same atom (e.g., `Man2S2P` is not possible).

### Compilers

In computer science, compilers translate source code into machine code. Source code is written in high-level, human-readable languages such as Java, C/C++, or Python. Machine code comprises a set of commands executed by the processor, such as adding the content of two registers or moving the content from one register to another. We apply the idea of compilers to the IUPAC notation. IUPAC annotations of molecules are more human-readable than SMILES, especially for molecules of the size of glycans that might contain hundreds of heavy atoms. Therefore, we treat IUPAC as a language and a specific, molecule-describing IUPAC string as a word of the new “IUPAC language”. These words are composed of so-called tokens that are combined following rules of the language, which jointly constitute a *grammar* [17].

Aside from the grammar, a compiler comprises a *lexer* and *parser* that performs the conversion from human-readable programming languages into machine code based on the grammar. Or, with GlyLES, from IUPAC words into SMILES. The lexer reads in the source code, in our case, the IUPAC language, and recognizes tokens in the input. Tokens are single or multiple characters, such as `def, lambda`, or: in Python. Examples from the IUPAC language for glycans are [or `Man`. The output of a lexer is a so-called token stream that is processed by a parser to generate a so-called *abstract syntax tree* (*AST*). The AST represents the internal structure of the input. Generating lexer and parser based on some grammar can be automated using tools such as ANTLR [18].

An example of this process can be seen when parsing (2 + 3) × 4 using simple arithmetic rules. These rules from analysis correspond to the parser rules of a compiler, whereas identifying the individual symbols as numbers, brackets, and arithmetic signs is what the lexer does. The output of the lexer, i.e. the token stream, is

~~~
(2 + 3) × 4
~~~

and the result of the parser, the AST, can be seen in Figure 1.

**Figure 1:**
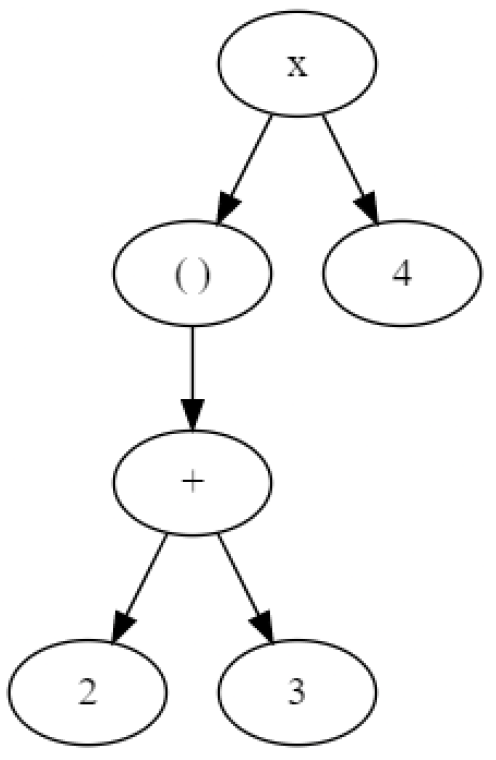
Example of an abstract syntax tree (AST) for the arithmetic expression (2 + 3) × 4.

**Figure 2:**
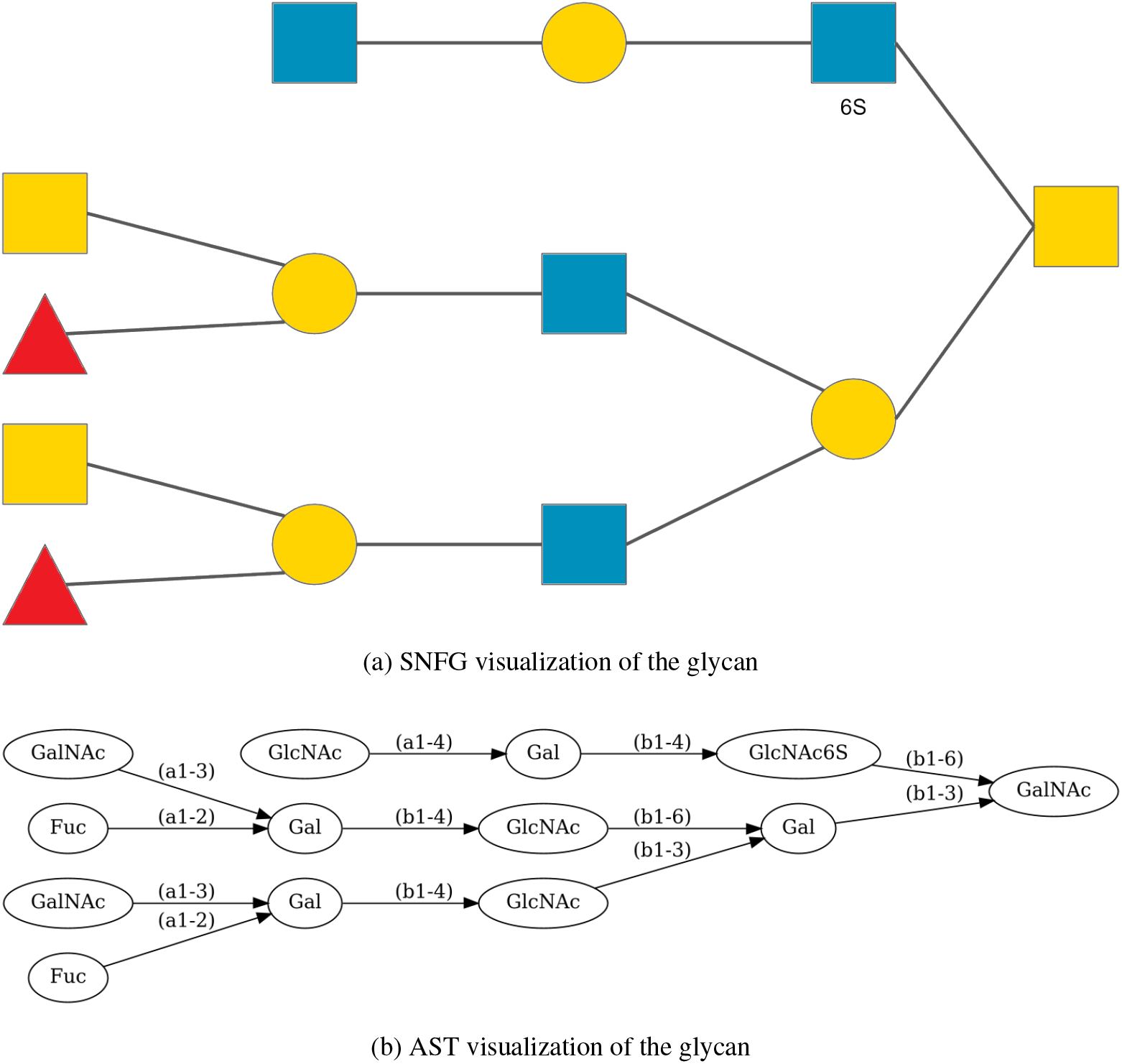
(a) SNFG image representation of the glycan in the implementation example. (b) AST representation of the glycan in the implementation example. In every node, we provide the IUPAC description for the single monosaccharide, including all its functional groups as the label.

### Previous work

To the best of our knowledge, there is no tool able to convert IUPAC notations of glycans into SMILES that works offline for arbitrary glycans in an automated fashion.

Alternatives can be divided into two groups: Databases that store information on glycans, including their SMILES string, from which mappings between IUPAC and SMILES strings can be extracted; and online tools converting IUPAC to SMILES.

Several databases, such as Glycopedia[19], GlyTouCan [20], and GlyGen [21], are specific for glycans. PubChem[22] also contains some glycans which are annotated with both IUPAC and SMILES. Yet all these databases are static and cannot be used to retrieve the SMILES notation for a not yet stored glycan, which is not uncommon given the large variety of glycan structures. Further, obtaining SMILES notation of a glycan motif can be cumbersome with this approach, as most databases only store full glycans, not motifs, and it is not human-readable which exact part of the SMILES string corresponds to which substructure in the IUPAC notation.

Among specialized online tools, REStLESS[23] and the “CSDB/SNFG structure editor”[24] are both based on the carbohydrates structure database[25]. REStLESS uses an IUPAC-condensed-like notation as a starting point when converting to SMILES. Neither of the tools is open-source or available for offline use. REStLESS may require pre-processing of input sequences, as they do not work on standard IUPAC-condensed glycans. Therefore, REStLESS cannot be seen as easy-to-use. The “CSDB/SNFG structure editor” has an online interface where one can build the glycan of choice and copy-paste the SMILES string, which is not workable to do in an automated fashion.

In this work, we present a tool that can convert arbitrary glycans from an IUPAC representation into SMILES representation that can further be used to build atomic graphs for the glycans.

### Implementation

The overall structure of the package comprises three major steps. First, we read in an IUPAC string and parse it into an AST based on a grammar. Second, we compute a SMILES string for each monomer in the nodes of the AST. The grammar for this can be found in the GitHub repository as well, https://github.com/kalininalab/GlyLES/blob/main/glyles/grammar/Glycan.g4 This includes adding all functional groups of the monosaccharide into a node. Finally, the tree is resolved into a single SMILES string representing the input IUPAC-encoded glycan in the SMILES notation. We will have a closer look at all these steps in the following sections.

### Step 1: Parsing IUPAC notation into AST

The IUPAC notation has a very regular structure to describe glycans. Knowing the internal structure of the IUPAC nomenclature for glycans, one can develop a grammar recognizing this general structure. Based on the grammar, there are tools such as ANTLR generating the lexer and parser for some grammar.

As described in section Compilers, the output of a parser is an AST, which is a tree of the tokens in the input. For a properly defined grammar, the AST is very similar to the structure of the glycan, as both structures are trees. Thus, the root monomer of the glycan corresponds to the root node of the AST. The leaf monosaccharides in the glycan can be found in the leaves of the AST. This has the benefit that we do not have to convert the output of the parser into some other structure. but can take it as it is.

For example, the AST for a glycan from human gastric mucins with an additional sulfate attached to a GlcNAc is very similar to the actual glycan structure (Figures 2a and 2b) [26]. The corresponding IUPAC string is

~~~
Fuc(a1-2)[
    GalNAc(a1-3)
]Gal(b1-4)GlcNAc(b1-3)[
    Fuc(a1-2)[
        GalNAc(a1-3)
    ]Gal(b1-4)GlcNAc(b1-6)
]Gal(b1-3)[
    GlcNAc(a1-4)Gal(b1-4)GlcNAc6S(b1-6)
]GalNAc.
~~~

We formatted the IUPAC with line-breaks wherever a new branch is introduced into the parent strand to better understand the structure.

### Step 2: Attaching functional groups

After parsing the IUPAC representation into an AST, all monomers of the glycan are stored in the nodes of the tree. Now, we have to compute the SMILES string for the monomer in each node, including all its modifications.

In this package, attaching functional groups to monosaccharides is done by the grammar using another helpful property of the IUPAC notation. From the example of `Gal(a1-6)3dManNAc4S(a1-4)Gal`, we can see that everything between the two bonds belongs to the mannose. Thus, the information on how to modify the SMILES description of the mannose can be stored directly in the node representing the mannose in the AST. The information on how to attach functional groups to monomers is provided in a rule in the grammar, and the different functional groups are given as tokens. This has the effect that every monosaccharide node in the AST stores information about the attached functional groups.

Once the AST is constructed, we can iterate over the nodes and generate a SMILES string for every monomer, including its functional groups. This is done by first identifying the monosaccharide in a node and whether it is in the alpha or beta enantiomeric form and creating its RDKit representation.To attach a functional group, we replace the atom in the monosaccharide the functional group binds to by a placeholder atom. In parallel, we look up the name of the functional group in a map from functional group names to SMILES strings. Then, the monosaccharide with the placeholder is converted to a SMILES string, the placeholder is replaced with the SMILES string of the functional group, and the SMILES is converted back into an RDKit molecule again.

In the example from the previous step, we zoomed in on a connection of two monosaccharides to show the procedure of step 2 in more detail (see Figure 3).

**Figure 3:**
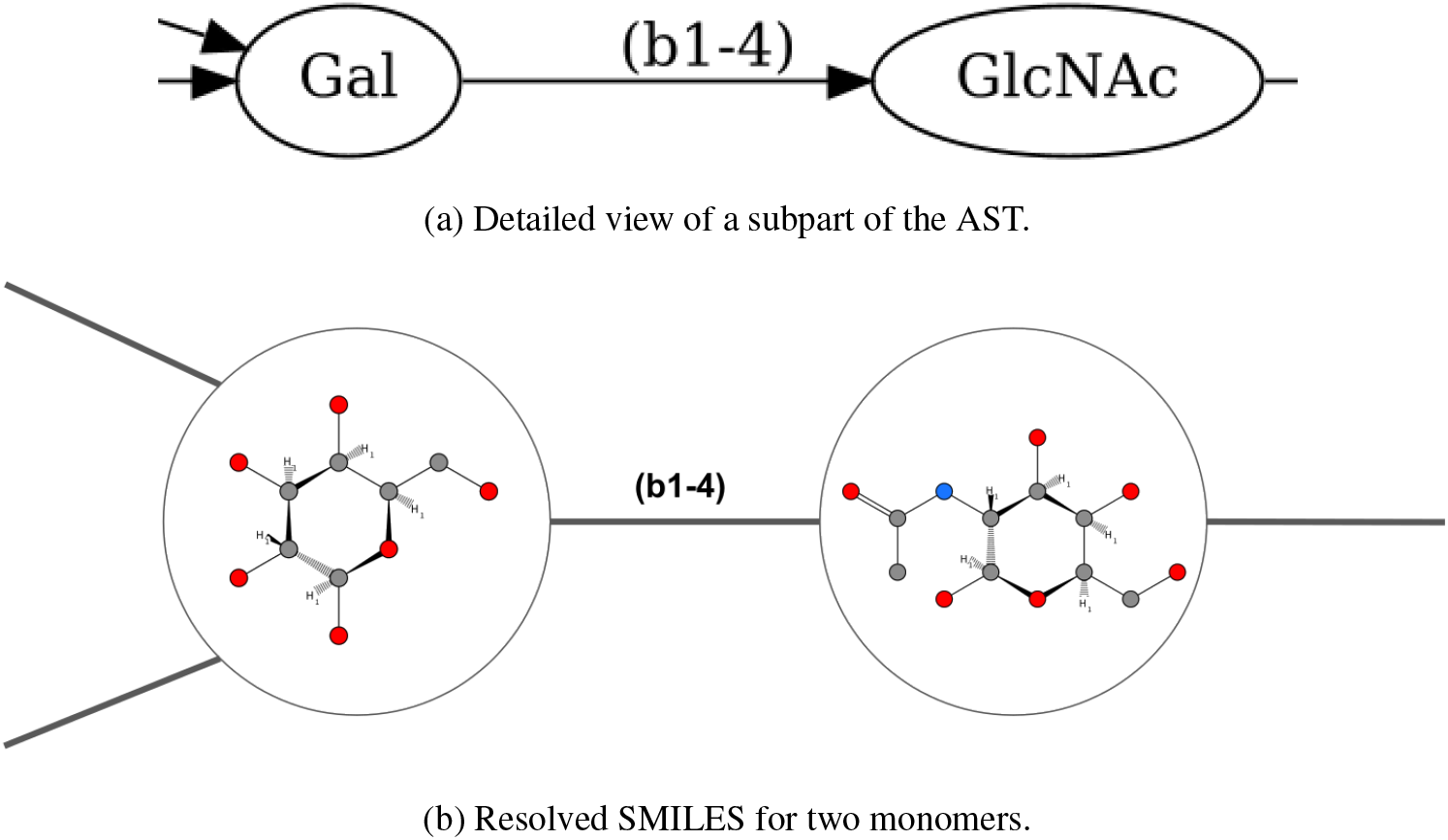
Adding functional groups. (a) Zoom into the AST displaying the branch between Gal and GlcNAc in the running example. (b) The corresponding part of the AST with monosaccharides resolved to atomic level and including modifications. Chemical structure visualization done with https://smarts.plus/

### Step 3: Connecting monomers

In the last step, we combine all the monomers in the nodes into a single SMILES string representing the entire glycan. This is done similarly to how we added the functional groups to the monosaccharides. First, we identify the position where a child monomer binds, replace that oxygen with a placeholder atom, generate a SMILES string and replace the placeholder atom with the SMILES string of the child monomer. This is done recursively until we put together the SMILES string for the root monosaccharide, which is the SMILES string for the represented glycan.

The result of this operation for the example in Figures 3a and 3b is

~~~
01[C@H](C0)[C@@H](
    Z
)[C@H](0)[C@@H](NC(C)=0)[C@@H]10
~~~

for the `GlcNAc b` where the SMILES string for `Gal b` will replace the `Z. Gal b` is translated into

~~~
0[C@@H]20[C@H](C0)[C@H](0)[C@H](
    X
)[C@H]2
    Y,
~~~

where `X` and `Y` will be replaced by the SMILES strings for `GalNAc a` and `Fuc` a from the parts of the tree that are not shown in figures 3a and 3b. If we plug in the part for `Gul a`, we get

~~~
01[C@H](C0)[C@@H](
    0[C@@H]20[C@H](C0)[C@H](0)[C@H](
        X
    )[C@H]2
        Y
)[C@H](0)[C@@H](NC(C)=0)[C@@H]10
~~~

showing the principle of how the SMILES string of the monosaccharides are plugged into each other.

## Results and Discussion

GlyLES has been tested using handcrafted tests aided by fuzzy tests using the pytest framework. Additionally, we scanned the entire PubChem database to collect a large set of examples to test if the tool works under real-world conditions. All these tests are included in the published code on GitHub (https://github.com/kalininalab/GlyLES). Comprehensive lists of all implemented monosaccharides and functional groups are provided as Supplementary Material.

The tool aims to convert any glycan with any combination of functional groups given by its IUPAC-condensed notation into a SMILES string. GlyLES can do this as a Python package that is easy to use, offline, and open source. Some rare monosaccharides and modifications may have not been implemented yet, but the tool will be updated as soon as they will be discovered.

For the glycans of the glycowork dataset [28] that have a complete structural description without wildcard connections between monosaccharides (~ 24, 000 structures), GlyLES can convert ~ 99%of the glycans. For this dataset, we have no labels and could only test if the output of the tool is a valid, organic molecule. We also extracted pairs of IUPAC-condensed and SMILES representations for glycan from the PubChem database to test if the tool also produces SMILES string representing the true molecules (~8,000 structures). In all cases, GlyLES reconstructed the correct atomic structure.

For example, the SMILES string for the molecule in figure 2a is

~~~
01C(0)[C@H](NC(C)=0)[C@@H](0[C@@H]20[C@H](C0[C@@H]30[C@H](C0)
[C@@H](0[C@@H]40[C@H](C0)[C@H](0)[C@H](0[C@H]50[C@H](C0)[C@H]
(0)[C@H](0)[C@H]5NC(C)=0)[C@H]40[C@@H]50[C@@H](C)[C@@H](0)[C@@H]
(0)[C@@H]50)[C@H](0)[C@H]3NC(C)=0)[C@H](0)[C@H](0[C@@H]30[C@H]
(C0)[C@@H](0[C@@H]40[C@H](C0)[C@H](0)[C@H](0[C@H]50[C@H](C0)
[C@H](0)[C@H](0)[C@H]5NC(C)=0)[C@H]40[C@@H]50[C@@H](C)[C@@H]
(0)[C@@H](0)[C@@H]50)[C@H](0)[C@H]3NC(C)=0)[C@H]20)[C@@H](0)
[C@H]1C0[C@@H]20[C@H](C0S(=0)(=0)0)[C@@H](0[C@@H]30[C@H](C0)
[C@H](0[C@H]40[C@H](C0)[C@@H](0)[C@H](0)[C@H]4NC(C)=0)[C@H]
(0)[C@H]30)[C@H](0)[C@H]2NC(C)=0.
~~~

Figure 4 visualizes this on an atomic level and highlights the single monomers with their symbols from SNFG.

**Figure 4:**
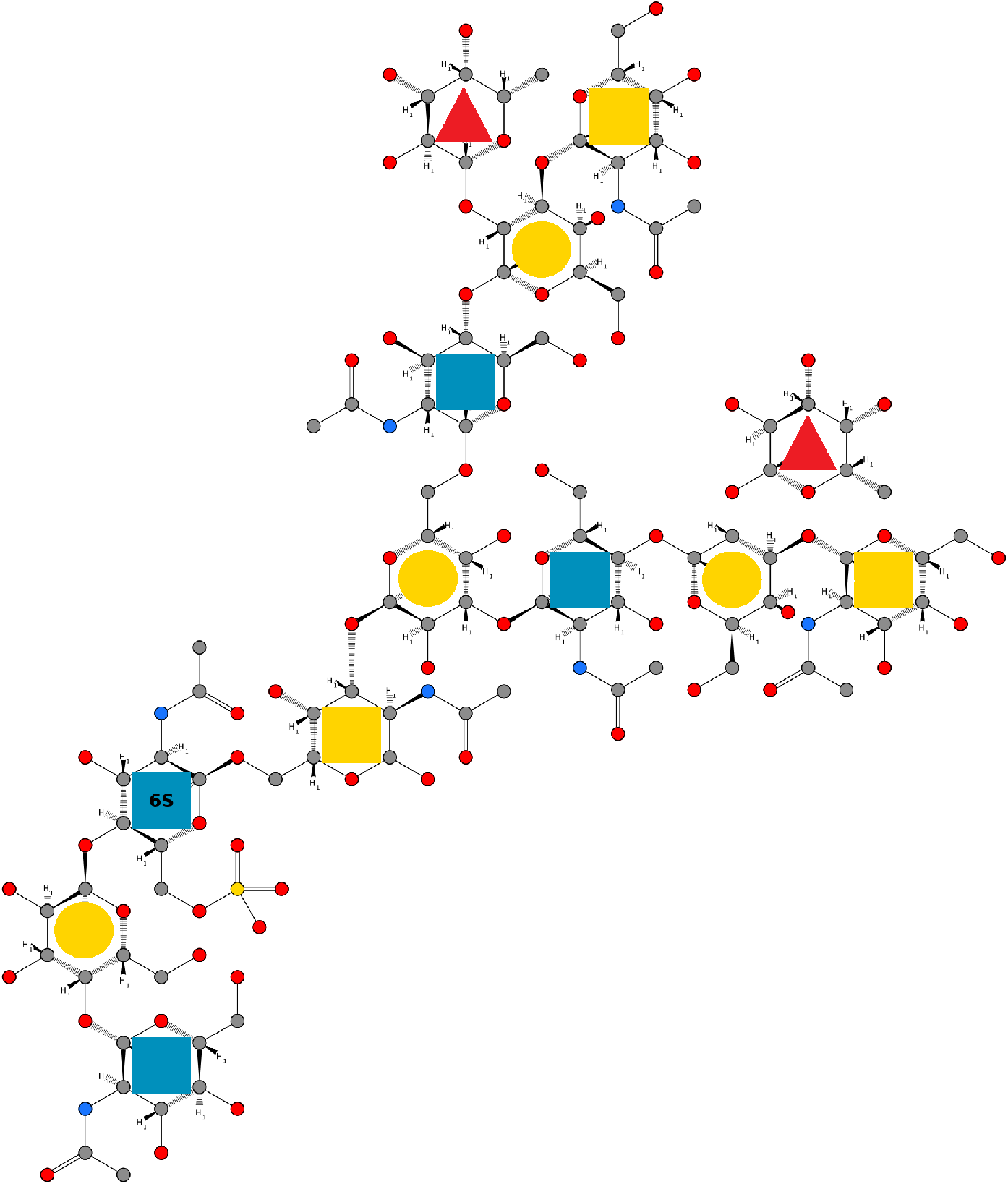
Visualization of the example glycan on an atomic level with the help of the GlyLES package. Chemical structure visualization done with https://smarts.plus/[27]

To estimate the average time that GlyLES needs to convert a glycan, we ran a timed test on a subset of our test glycans. We took those glycans we had the SMILES strings for, to be sure GlyLES does not raise an error and the output will be correct. Error-free output is important because raising an error costs additional time and would distort the result. The tested glycans differ largely in their size and shape to get a better averaged estimate of the runtime. Overall, we converted 11,951 glycans into a list of SMILES strings. This process lasts ~232 seconds, which results in an average time per glycan of ~19 milliseconds. This measurement was conducted on a single 11th Gen Intel Core i7-11700 with 2.5 GHz.

## Conclusion

To our knowledge, GlyLES is the first free, offline, and open-source tool to convert glycan representations from IUPAC-condensed notation to SMILES. It can compute SMILES representations of glycans in research projects where glycans need to be represented on an atomic level. Such applications include, but are not limited to, glycan visualization, substructure search and comparison, molecular modelling, docking and molecular dynamics, or featurization for machine learning and deep learning models. GlyLES is more flexible, extendable, and easier to maintain than databases which are primarily used so far. It is also more generalizable and efficient than databases, as all glycans are computed during runtime and it is not necessary to store anything.

One limitation of GlyLES is that IUPAC allows for wildcards at positions where the structure of a glycan is not fully resolved. For example, for a string `Man(?1-?)Gal` it is known that a mannose and a galactose are connected, but it is not resolved how. In fact, there are 10 distinct possibilities, how to resolve this wildcard bond. This is possible in IUPAC but not in SMILES. Therefore, GlyLES cannot convert such structures.

## Availability and Requirements

**Project Name:** GlyLES

**Project home page**: https://github.com/kalininalab/GlyLES

**Operating system(s)** Platform independent

**Programming language:** Python

**Other requirements:** None

**License:** MIT License

**Any restriction to use by non-academics:** No restrictions

## Declarations

## Acknowledgements

The authors want to thank Alexander Titz for fruitful discussions about glycans and their structures.

## Funding

R.J. was partially funded by the HelmholtzAI project XAI-Graph. D.B. was funded by a Branco Weiss Fellowship – Society in Science, the Knut and Alice Wallenberg Foundation, and the University of Gothenburg, Sweden. O.V.K. acknowledges financial support from the Klaus-Faber Foundation.

## Abbreviations

ANTLR: Another Tool for Language Recognition
ASCII: American Standard Code for Information Interchange
AST: abstract syntax tree
Gal: Galactose
GalNAc: N-Acetyl-Galactosamine
GlcNAc: N-Acetyl-Glucosamine
IUPAC: International Union of Pure and Applied Chemistry
MAL-1: Maackia amurensis-I
Man: Mannose
MurNAc: N-Acetyl Muramic Acid
Neu5Ac: N-Acetyl Neuraminic Acid
SMILES: Simplified Molecular-Input Line-Entry System)
SNFG: Symbol Nomenclature for Glycans
WGA: Wheat germ agglutinin

## Availability of data and materials

Archived version under the project home page, see the Availability and Requirements section for further details.

## Ethics approval and consent to participate

Not applicable

## Competing interests

The authors declare that they have no competing interests.

## Consent for publication

Not applicable

## Authors’ contributions

R.J. implemented and tested the code. D.B. and O.V.K. helped with the documentation and the structures of glycans. All authors read and approved the final manuscript.

## Notes

### Competing Interest Statement

The authors have declared no competing interest.

